# Meningeal CSF transport is primarily mediated by the arachnoid and pia maters during development

**DOI:** 10.1101/2023.08.10.552826

**Authors:** Shelei Pan, Joshua Koleske, Gretchen M. Koller, Grace L. Halupnik, Abdul-Haq O. Alli, Shriya Koneru, Dakota DeFreitas, uthi Ramagiri, Jennifer M. Strahle

## Abstract

**Background:** The recent characterization of the glymphatic system and meningeal lymphatics has re-emphasized the role of the meninges in facilitating CSF transport and clearance. Here, we characterize small and large CSF solute distribution patterns along the intracranial and surface meninges in neonatal rodents and compare our findings to a rodent model of intraventricular hemorrhage-posthemorrhagic hydrocephalus. We also examine CSF interactions with the tela choroidea and its pial invaginations into the choroid plexuses of the lateral, third, and fourth ventricles.

**Methods:** 1.9-nm gold nanoparticles, 15-nm gold nanoparticles, or 3 kDa Red Dextran Tetramethylrhodamine constituted in aCSF were infused into the right lateral ventricle of P7 rats to track CSF circulation. 10 minutes post-1.9-nm gold nanoparticle and Red Dextran Tetramethylrhodamine injection and 4 hours post-15-nm gold nanoparticle injection, animals were sacrificed and brains harvested for histologic analysis to identify CSF tracer localization in the cranial and spine meninges and choroid plexus. Spinal dura and leptomeninges (arachnoid and pia) wholemounts were also performed.

**Results:** There was significantly less CSF tracer distribution in the dura compared to the arachnoid and pia maters in neonatal rodents. Both small and large CSF tracers were transported intracranially to the arachnoid and pia mater of the perimesencephalic cisterns and tela choroidea, but not the dura mater of the falx cerebri. CSF tracers followed a similar distribution pattern in the spinal meninges. In the choroid plexus, there was large CSF tracer distribution in the apical surface of epithelial cells, and small CSF tracer along the basolateral surface. There were no significant differences in tracer intensity in the intracranial meninges of control vs. intraventricular hemorrhage-posthemorrhagic hydrocephalus (PHH) rodents, indicating preserved meningeal transport in the setting of PHH.

**Conclusions:** Differential CSF tracer handling by the leptomeninges suggests that there are distinct roles for CSF handling between the arachnoid-pia and dura maters in the developing brain. Similarly, differences in apical vs. luminal choroid plexus CSF handling may provide insight into particle-size dependent CSF transport at the CSF-choroid plexus border.

## Introduction

A significant challenge to the study of cerebrospinal fluid (CSF) is the relative lack of understanding for how CSF (and its solutes) circulate within and out of the central nervous system (CNS). The canonical theory of CSF circulation posits that CSF is produced in the choroid plexuses (ChP) of the ventricles before flowing from the ventricular system to the subarachnoid space (SAS) and basal cisterns, where it exits the brain through the venous system by way of arachnoid villi and granulations [1]. However, arachnoid villi and granulations do not mature until two years of age in humans and are entirely absent or non-functional in various mammalian species including rodents [2–5]. These findings do not support arachnoid villi-mediated CSF outflow and suggest that alternative routes exist for CSF circulation and efflux.

Two candidate theories for CSF clearance that have emerged in recent years are the glymphatic theory and meningeal lymphatics [6,7]. The reinvigorated focus on the meninges heralded by these two discoveries is notable considering the role of the meninges in CSF circulation has historically been the subject of longstanding speculation. 19^th^ century French physiologist Francois Magendie, who discovered the eponymous foramen of Magendie, postulated that the leptomeninges (pia and arachnoid mater) produce CSF [1,8]. Even though this theory was later disregarded in favor of histologic, pharmacologic, and physiologic studies supporting the ChP as the cite of CSF production [9–14], his discoveries highlighted the notion that the meninges are more than a protective barrier. It is now known that the meningeal lymphatics in the dorsal convexities and at the base of the skull play a role in CSF solute drainage [7,15], and that the leptomeninges and dura mater have different transcriptional signatures and play distinct roles within the CNS [16]. In addition to their spatial heterogeneity, meningeal fibroblasts also have distinct transcriptional signatures across development [16], however it is not clear if meningeal CSF solute handling during development differs from what is observed in adult and aged animals. Additionally, it is not clear if there are differences in CSF handling within the layers of the neonatal meninges including the arachnoid and pia (vs. dura). As the relationship between meninges and CSF becomes increasingly intertwined yet complex, it is important to understand where in the meninges CSF is handled prior to its clearance from the brain.

In this present study, we present a CNS-wide characterization of CSF solute movement within the meninges and its related structures in neonatal rodents, with special emphasis on the intracranial meninges, spinal meninges, and ChP. Using intraventricular injections of large (15-nm) and small (1.9-nm and 3 kDa) CSF tracers, we show that both large and small CSF tracers distribute primarily within the cranial and spine leptomeninges with more limited distribution within the dura. Both large and small CSF tracers circulate through the leptomeninges of the perimesencephalic cisterns and are also present in the third ventricular tela choroidea and velum interpositum, a distribution pattern that is not affected by intraventricular hemorrhage- posthemorrhagic hydrocephalus. In the ChP, small CSF tracers are found in the luminal pial membrane and the basolateral surface of the choroid plexus epithelial cells, while large CSF tracers preferentially accumulate on the apical surface. Finally, meningeal transport is preserved early after PHH and may act as a route to transmit both physiologic CSF solutes and blood breakdown products after intraventricular hemorrhage.

## Methods

### Animals (Rodents)

All experiments were approved by the Institutional Animal Care and Use Committee of Washington University (protocol #22-0614). Sprague Dawley Rats (crl:SD400, Charles River Laboratories, Wilmington, MA) were used in all experiments. Female and male post-natal day 4-7 Sprague Dawley Rats were housed with their dams in a 12-hour light-dark cycle in a temperature and humidity-controlled room. Water and food were provided ad libitum for the dam.

### Intraventricular hemorrhage-posthemorrhagic hydrocephalus induction

P4 rodents were anesthetized (isoflurane 2-3% induction and 1.5% maintenance) and fixed in a stereotaxic frame. A 2.5 mm midline incision was made and a 0.3cc syringe with a 30-gauge needle was inserted into the right lateral ventricle at the following coordinates from bregma: 1.5 mm lateral, 0.4 mm anterior, and 2.0 mm deep. A small volume (20 μL) of artificial CSF (aCSF) (Tocris Bioscience, Bristol, UK) or hemoglobin constituted in aCSF at a 150 mg/mL concentration was injected at a rate of 8000 nL/min using a micro-infusion pump (World Precision Instruments, Sarasota, FL) to create the aCSF control and IVH-PHH conditions respectively. The needle was left in for 5 minutes post injection to prevent backflow. The incision was closed with 6-0 Ethilon suture (Ethicon Inc, Raritan, NJ). Rodents recovered from anesthesia and were returned to their cage with the dam for 72 hours before CSF tracer injection at P7.

### CSF tracer injections

Anesthetized naïve (Figure 1B-D, 2A, 4B-D, 5A, 8), aCSF control (Figure 1F, 2B-F, 3, 4E, 5B-I, 6, 7, Supplementary Figure 2), and IVH-PHH (Supplementary Figure 2) P7 rats underwent intraventricular injection of 20 µl of 3 kDa Red Dextran Tetramethyl rhodamine (RD/TMR) (D3307, Thermo Fisher Scientific, Waltham, MA), 1.9-nm (1102, Nanoprobes, Yaphank, NY), or 15-nm (1115, Nanoprobes, Yaphank, NY) as per the protocol above with the following coordinates from bregma: 1.7 mm lateral, 0.5 mm anterior, and 2.0 mm deep. 1.9- and 15-nm AuNPs were constituted in aCSF at a concentration of 200 mg/mL, while RD/TMR was dissolved in aCSF in a 0.25% w/v solution. Rodents were sacrificed with intracardial perfusion with 10 mL of ice-cold PBS followed by 10 mL of 4% PFA at 10 minutes after 1.9-nm AuNP and 3 kDa RD/TMR injections or 4 hours after 15-nm AuNP injections.

**Figure 1.**
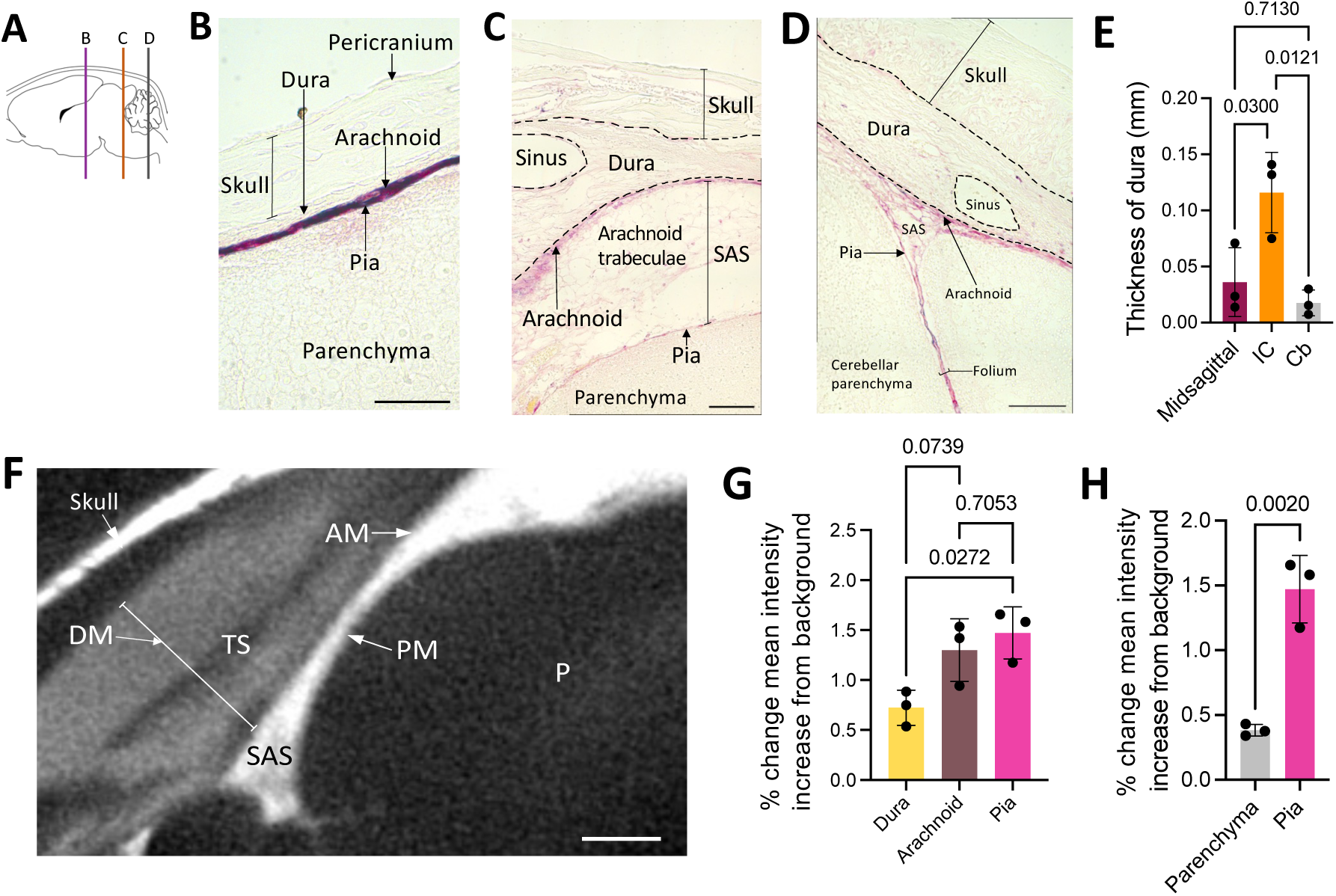
Large CSF tracers primarily circulate within the neonatal arachnoid and pia maters and not the dura. **A**, Schematic of regions shown in B, C, and D. **B-D**, Representative histology of decalcified skulls and the underlying meninges and parenchyma over the cortex (B), inferior colliculus (C), and cerebellum (D) showing minimal large CSF tracer (magenta) distribution through the pericranium, skull, and dura mater, but widespread distribution within the arachnoid and pia maters 4 hours after 15-nm gold nanoparticle (AuNP) injection into the right lateral ventricle of P7 rats. There was also limited AuNP influx into the parenchyma. The subarachnoid space is collapsed post-mortem and differentiation between layers was primarily based on tissue morphology. scalebars = 50 µm. **E,** Quantification of the thickness of the midsagittal dura over the longitudinal fissure, the inferior colliculus (IC), and cerebellum (Cb). Data are mean ± SD, n = 3 per group; One-way ANOVA with post-hoc Tukey. **F,** Representative X-ray microtomography (XRM) image showing high amounts of AuNP enhancement within the arachnoid mater (AM), pia mater (PM), and subarachnoid space (SAS), with less enhancement in the dura mater (DM). There was minimal enhancement in the lumen of the transverse sinus (TS) and parenchyma (P). scalebar = 500 µm. **G-H,** Quantification of the percent change in mean intensity increase of 15-nm AuNPs in the dura, arachnoid, and pia maters (G) and the pia mater and parenchyma (H) compared to the background XRM signal. Data are mean ± SD, n = 3 per group; G, One-way ANOVA with post-hoc Tukey. H, Unpaired, two-tailed t-test All data are representative of 3 rodents.

**Figure 2.**
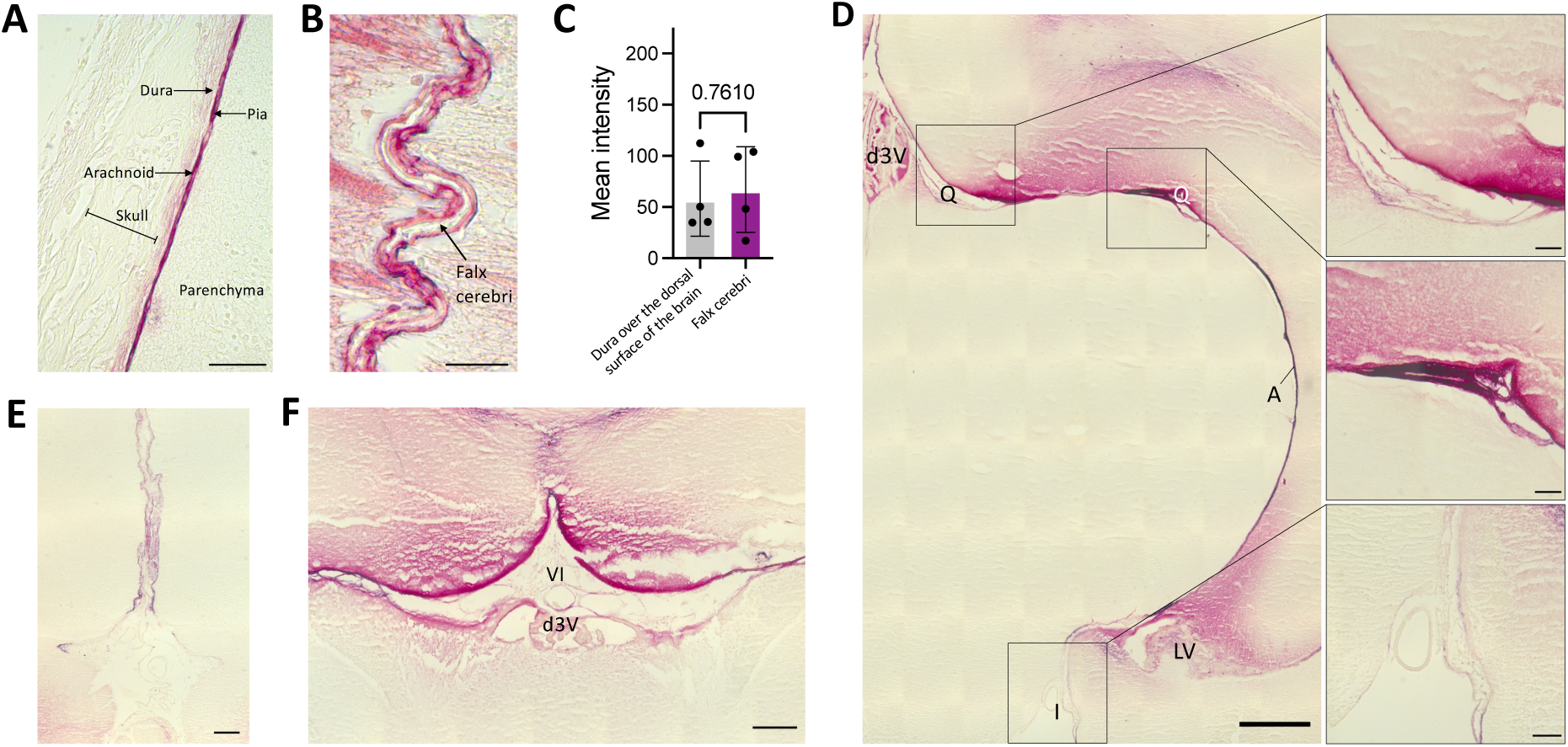
Distribution of large CSF tracers within the intracranial meninges. **A,** Representative histology of the meninges (including the dura, arachnoid, and pia maters) and the overlying decalcified skull 4 hours after 15-nm gold nanoparticle (AuNP) injection into the right lateral ventricle of P7 rats. scalebar = 50 µm. **B,** Representative histology showing 15-nm AuNP distribution within the falx cerebri and leptomeningeal (pia and arachnoid) invaginations into the longitudinal fissure. The falx cerebri is labeled with a black arrow. scalebar = 25 µm. **C,** Quantification of the mean intensity of 15-nm AuNP distribution within the dura mater over the surface of the brain compared to the falx cerebri. Data are mean ± SD, n = 3 per group; Unpaired, two-tailed t-test. **D,** Representative histology showing 15-nm AuNP circulation through the dorsal third ventricle (d3V), quadrigeminal cistern (Q), ambient cistern (A), interpeduncular cistern (I), and lateral ventricle (LV) 4 hours after intraventricular injection in P7 rodents. d scalebar = 500 µm, d inset scalebars = 75 µm. **E-F,** 15-nm AuNP distribution in the rhinal fissure (E) and velum interpositum (F). Abbreviations: VI, velum interpositum; d3V, dorsal third ventricle. e scalebar = 100 µm, f scalebar = 250 µm. All data are representative of 3 rodents.

**Figure 3.**
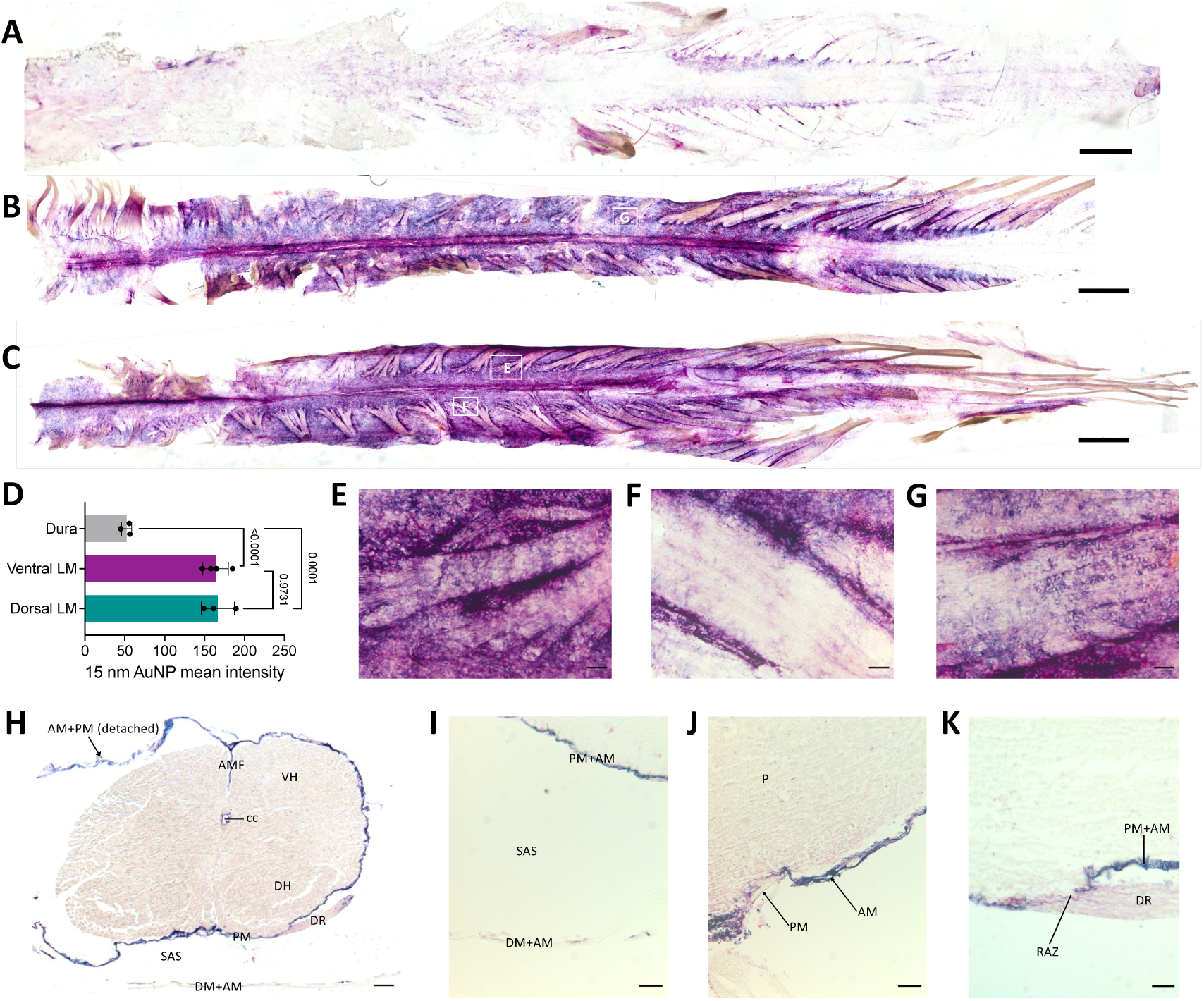
Large CSF tracers preferentially circulate within the spinal leptomeninges with limited entry superiorly into the spinal dura and inferiorly into the underlying parenchyma. **A-B,** Representative wholemounts showing large CSF tracer distribution through the dorsal spinal cord dura (A) and dorsal spinal cord leptomeninges (pia and arachnoid) (B) 4 hours after 15-nm gold nanoparticle (AuNP) injection into the right lateral ventricle of P7 rodents. scalebar = 1 cm. **C,** Representative wholemount showing 15-nm AuNP distribution through the ventral spinal cord leptomeninges 4 hours after intraventricular injection into the right lateral ventricle of P7 rodents. scalebar = 1 cm. **D**, Quantification of 15-nm AuNP mean intensity in the spine dura, ventral leptomeninges, and dorsal leptomeninges. There was significantly more 15-nm AuNP distribution in the leptomeninges than the dura. Data are mean ± SD, n = 3 per group; One-way ANOVA with post-hoc Tukey. **E-G,** High magnification histology of 15-nm AuNP distribution along the leptomeninges of spinal nerve roots as they leave the spinal cord. 15- nm AuNPs distributed primarily around the roots (G), with additional diffuse 15-nm AuNPs observed inside the roots (E, F). Areas from which high-magnification images are obtained are indicated by white boxes in 2B and 2C. E-G scalebars = 25 µm. **H-K,** Representative cross-sectional histology showing 15-nm AuNP distribution in the spinal cord, spinal leptomeninges and spinal dura mater. The anterior median fissure (AMF), aqueduct (Aq), dorsal horn (DH), dorsal root (DR), dura mater (DM), arachnoid mater (AM), pia mater (PM), ventral horn (VH), subarachnoid space (SAS), root attachment zone (RAZ), and parenchyma (P) are indicated. H scalebar = 1 mm, I-K scalebars = 25 µm. All data are representative of 3 rodents.

**Figure 4.**
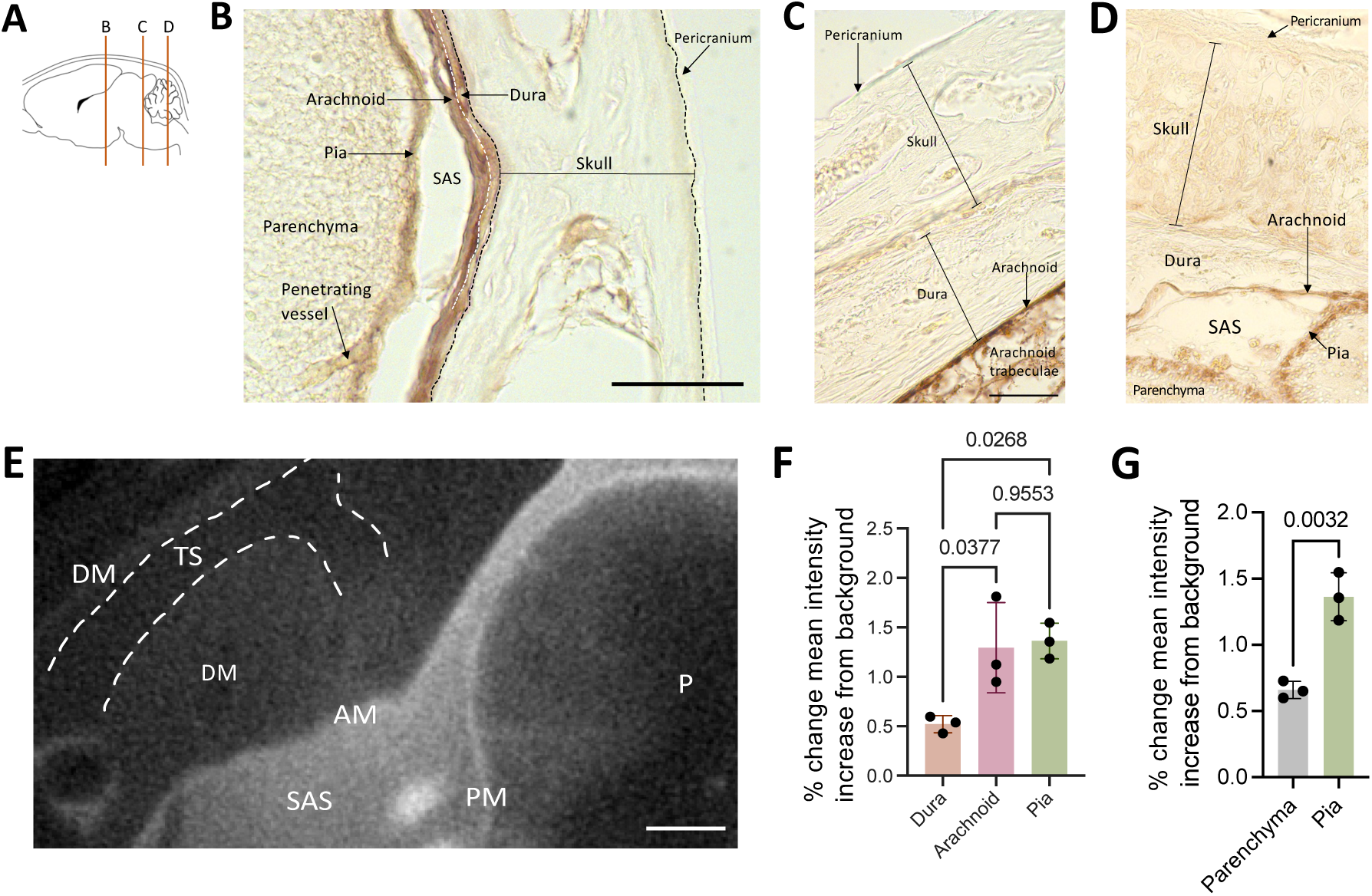
Small CSF tracers primarily circulate within the leptomeninges and not the dura. **A,** Schematic of regions shown in B, C, and D. **B-D**, Representative histology of decalcified skulls and the underlying meninges and parenchyma showing widespread small CSF tracer distribution within the arachnoid and pia maters over the cortex (B), inferior colliculus (C), and cerebellum (D) 10 minutes after 1.9-nm gold nanoparticle (AuNP) injection into the right lateral ventricle of P7 rats. There was minimal 1.9-nm AuNP (brown) distribution through the pericranium and skull, and limited tracer distribution through the dura mater along the surface of the brain. The subarachnoid space is collapsed post-mortem and differentiation between layers was primarily based on tissue morphology. scalebar = 50 µm. **E,** Representative X-ray microtomograph (XRM) showing 1.9-nm AuNP distribution within the pia mater (PM), arachnoid mater (AM), subarachnoid space (SAS), and parenchyma (P) at the level of the inferior colliculus. Similar to observations on histology, there was minimal enhancement in the dura mater (DM) and little to no enhancement in the transverse sinus (TS, dashed white line). scalebar = 500 µm. **F-G,** Quantification of the percent change in mean intensity increase of 1.9-nm AuNPs in the dura, arachnoid, and pia maters (F) and pia mater and parenchyma (G) compared to the background XRM signal. Data are mean ± SD, n = 3 per group; F, One-way ANOVA with post-hoc Tukey. G, unpaired, two-tailed T test. All data are representative of 3 rodents.

**Figure 5.**
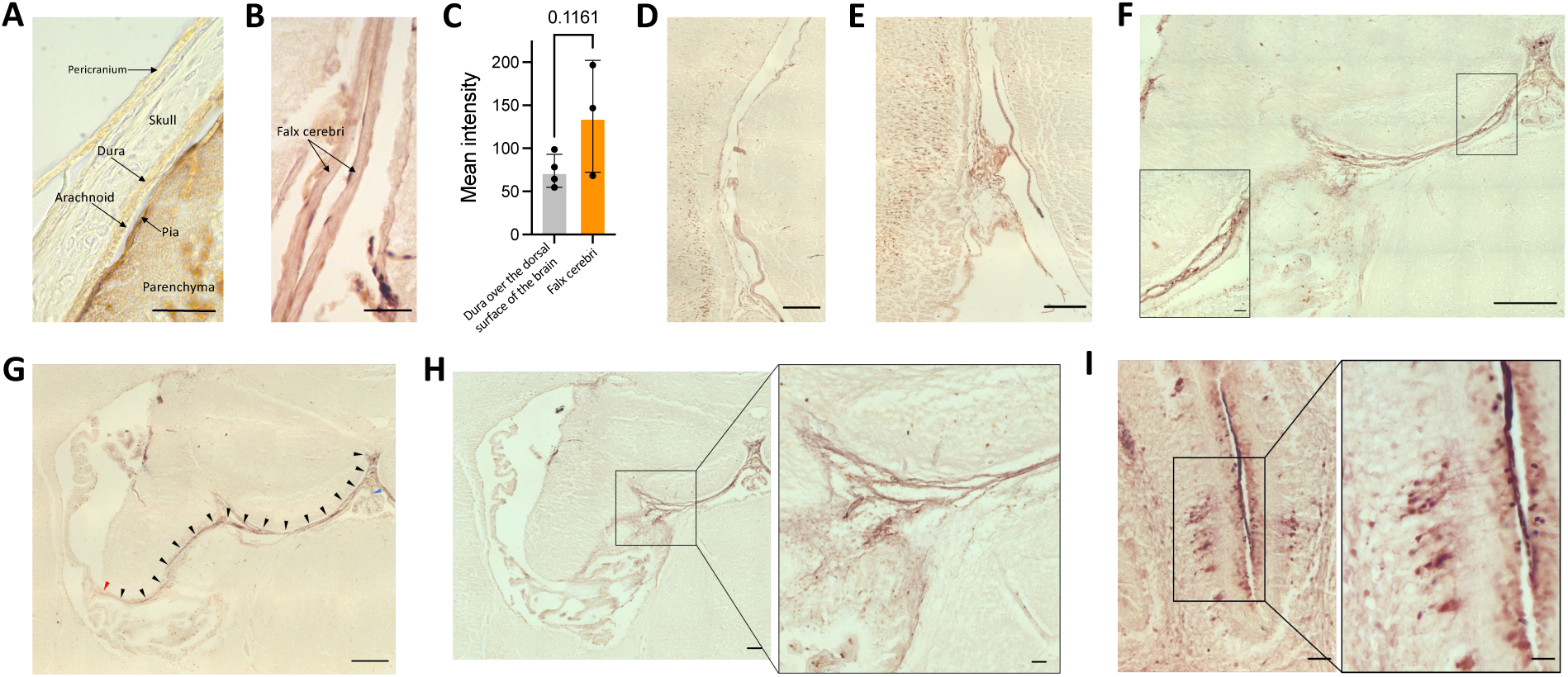
Distribution of small CSF tracers within the intracranial meninges. **A,** Representative histology showing small CSF tracer (brown) distribution through the decalcified skull, underlying meninges (dura, arachnoid, and pia maters), and parenchyma 10 minutes after 1.9-nm gold nanoparticle (AuNP) injection into the right lateral ventricle of P7 rodents. scalebar = 50 µm. **B,** Representative histology showing 1.9- nm AuNP distribution within the meningeal invaginations into the longitudinal fissure. The falx cerebri is labeled with a black arrow. scalebar = 50 µm. B scalebar = 100 µm. **C,** Quantification of the mean intensity of 1.9-nm AuNP distribution within the dura mater over the surface of the brain compared to the falx cerebri. Data are mean ± SD, n = 3 per group; Unpaired, two-tailed t-test. **D-F,** Representative histology showing 1.9-nm AuNP distribution within the intracranial meninges of the ambient cistern (D), interpeduncular cistern (E), and choroidal fissure (F). D scalebar = 250 µm, E scalebar = 250 µm, F scalebar = 250 µm. **G-H**, Representative histology showing the path the tela choroidea takes through the choroidal fissure from the roof of the third ventricle to the choroid plexus of the lateral ventricle (F) (black arrowheads), and the tela choroidea pia mater as it gives rise to the right lateral ventricle choroid plexus (G, H) (red arrowhead, G; inset H). G scalebar = 250 µm; H scalebar = 100 µm, H inset scalebar = 50 µm. **I,** Representative histology of 1.9-nm AuNP distribution in the leptomeningeal invaginations into the folia of the cerebellum. I scalebar = 25 µm, I inset scalebar = 10 µm. All data are representative of 3 rodents.

**Figure 6.**
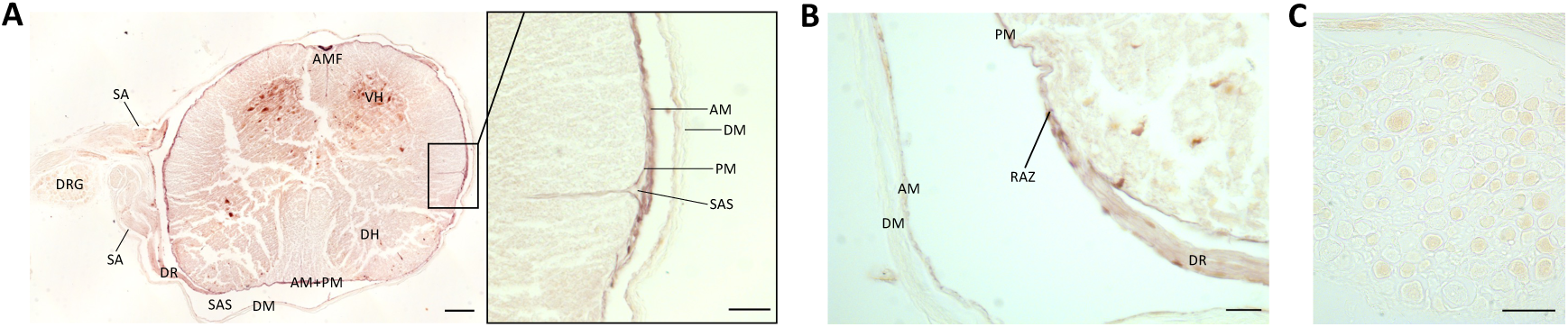
Small CSF tracers predominantly circulate within the leptomeninges of the spinal cord and parenchyma. **A-B,** Cross sectional histology showing small CSF tracer distribution (brown) in the spinal cord parenchyma, spine leptomeninges, and spine dura 10 minutes after 1.9-nm gold nanoparticle (AuNP) injection into the right lateral ventricle in P7 rodents. The anterior median fissure (AMF), ventral horn (VH), subarachnoid angle (SA), dorsal root ganglion (DRG), dorsal root (DR), pia mater (PM), subarachnoid space (SAS), dura mater (DM), root attachment zone (RAZ), and arachnoid mater (AM) are indicated. A scalebar = 1 mm, A inset scalebar = 25 µm, B scalebar = 25 µm. **C,** High magnification histology showing 1.9-nm AuNP distribution in the dorsal root ganglion. scalebar = 50 µm. All data are representative of 3 rodents.

**Figure 7.**
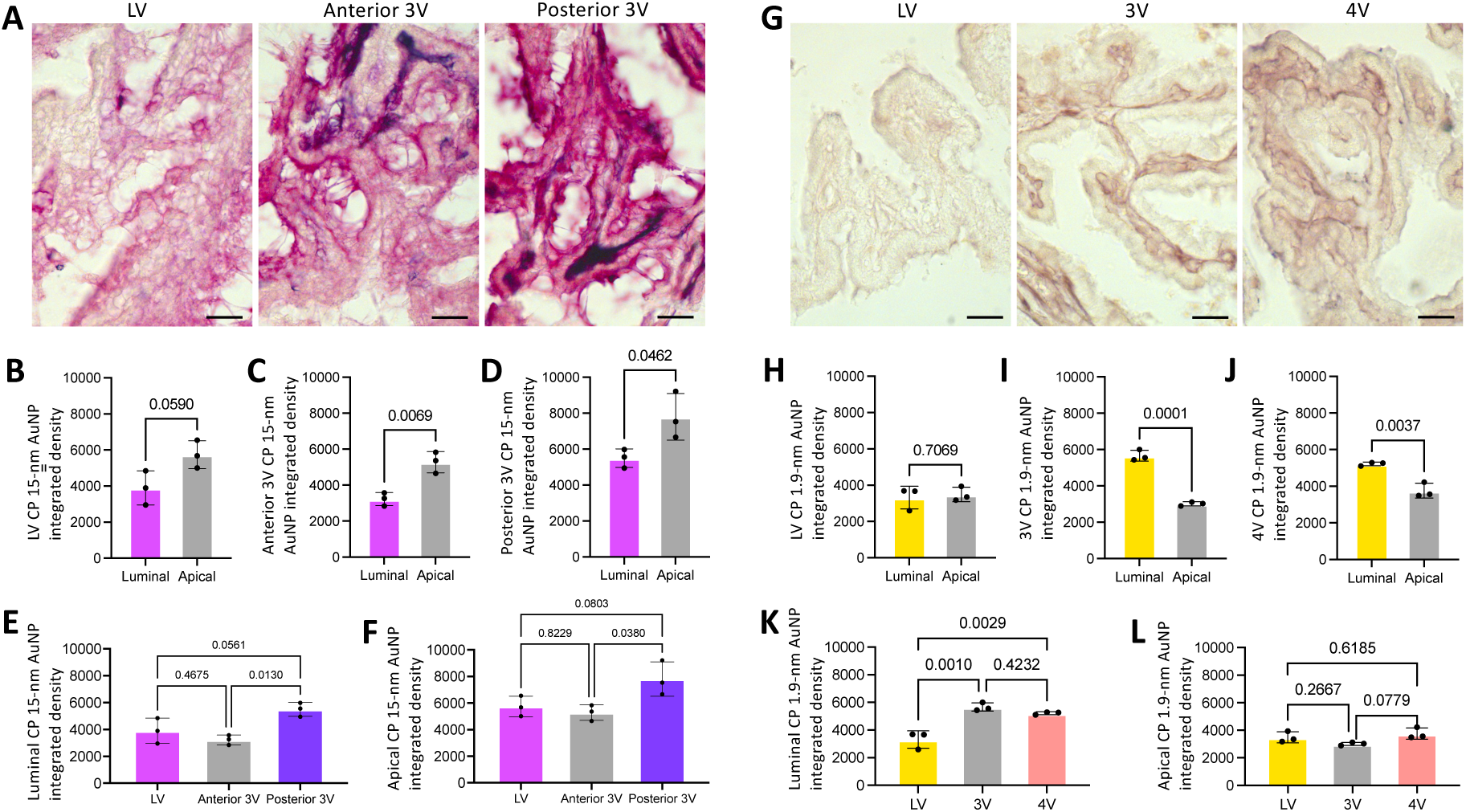
CSF tracer distribution within the choroid plexus. **A,** Representative histology of large CSF tracer distribution in the lateral ventricle (LV), anterior third ventricle (3V), and posterior 3V choroid plexuses (ChP) 4 hours post-15-nm gold nanoparticle (AuNP) injection into the right lateral ventricle of P7 rodents. scalebars = 25 µm. **B-D,** Quantification of 15-nm AuNP integrated density along the luminal and apical surfaces of the LV (B), anterior 3V (C), and posterior 3V (D) ChP. 15-nm AuNPs distributed along the apical surface of the anterior and posterior 3V ChP significantly more than the luminal surface. Data are mean ± SD, n = 3 per group; Unpaired two-tailed t test. **E-F,** Quantification of 15-nm AuNP distribution in the LV, anterior 3V, and posterior 3V ChP along the luminal (E) and apical (F) surfaces. There was significantly more 15-nm AuNP distribution on both the luminal and apical surfaces of the posterior 3V ChP compared to the anterior 3V ChP (F). Data are mean ± SD, n = 3 per group; One-way ANOVA with post-hoc Tukey. All data are representative of 3 rodents. **G,** Representative photomicrographs of small CSF tracer (brown) distribution in the lateral ventricle (LV), third ventricle (3V), and fourth ventricle (4V) choroid plexuses (ChP) 10 minutes post-1.9-nm gold nanoparticle (AuNP) injection into the lateral ventricle in P7 rodents. scalebars = 25 µm. **H-J,** Quantification of 1.9-nm AuNP integrated density between the luminal and apical surfaces of the LV (H), 3V (I), and 4V (J) ChP. There was significantly more AuNP distribution along the luminal surface of the 3V and 4V ChP compared to the apical surface. Data are mean ± SD, n = 3 per group; Unpaired two-tailed t test. **K-L,** Quantification and comparison of 1.9-nm AuNP distribution between the LV, 3V, and 4V ChP along the luminal (K) and apical surfaces (L). There was significantly more 1.9-nm AuNP distribution throughout the luminal 3V and 4V ChP compared to the LV, but no differences between regions on the apical ChP. Data are mean ± SD, n = 3 per group; One-way ANOVA with post-hoc Tukey. All data are representative of 3 rodents.

**Figure 8.**
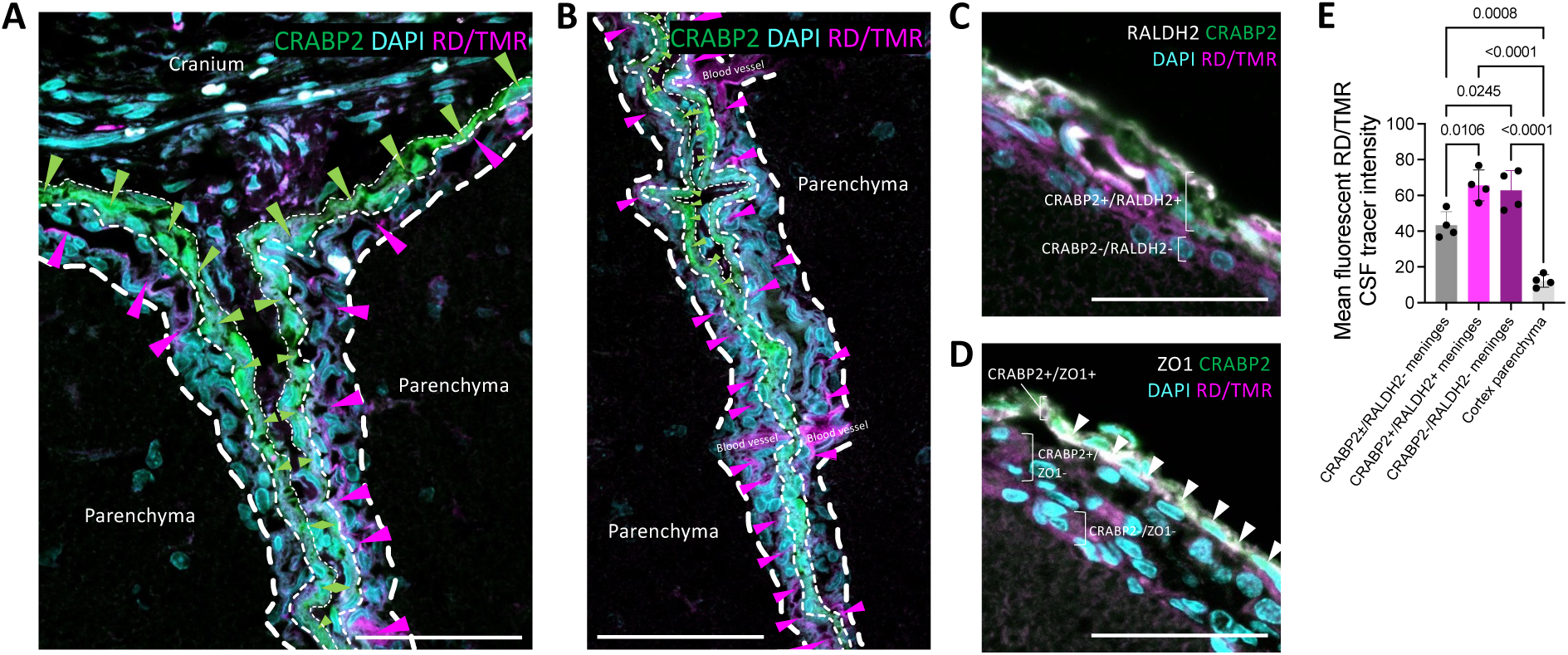
Immunofluorescent co-localization of small fluorescent CSF tracers with meningeal fibroblast markers. **A-B**, Representative photomicrograph of 3 kDa Red Dextran Tetramethylrhodamine (RD/TMR) (magenta, magenta arrowheads) distribution along the meninges around (A) and within (A, B) the longitudinal fissure 10 minutes after intraventricular injection into the right lateral ventricle in P7 rodents. RD/TMR distribution is primarily within the meningeal layer(s) inferior to a cellular retinoic acid binding protein 2 (CRABP2) (green, green arrowheads)-positive layer (inferior layer identified as the pia mater), with additional RD/TMR co-localization with the CRABP2+ layer (identified as the arachnoid mater). The putative border between the pia mater and underlying parenchyma is indicated with thick white dashed line; the pia and arachnoid are separated by the medium white dashed line, and the arachnoid and dura (falx) are separated by the thin white dashed line. Very little RD/TMR is seen medial/superior to the CRABP2+ layer (dura mater). A-B scalebars = 50 µm. **C-D**, Representative photomicrographs of RD/TMR distribution in the meninges over the dorsal surface of the brain and CRABP2, retinaldehyde dehydrogenase 2 (RALDH2), and zonula occludens 1 (ZO-1) staining. RD/TMR co- localized with a CRABP2+/RALDH2+ layer (arachnoid mater), in addition to the inferiorly adjacent CRABP2-/RALDH2- layer (pia mater) (C). RD/TMR distribution was inferior to the CRABP2+/ZO-1+ layer (arachnoid barrier cell layer), suggesting it localizes to the arachnoid and pia maters (D). **E**, Quantification of RD/TMR mean intensity in CRABP2±/RALDH2- meninges (dura), CRABP2+/RALDH2+ meninges (arachnoid), CRABP2-/RALDH2- meninges (pia), and the parenchyma of the cortex. There was significantly less RD/TMR in the dura and cortex parenchyma compared to the pia and arachnoid. Data are mean ± SD, n = 4 per group; One-way ANOVA with post-hoc Tukey. All data are representative of 4 rodents.

#### Sample preparation for meninges analysis using light microscopy

After intracardial perfusion and sacrifice, the cranium was harvested whole in a subset of rodents (Figure 1B-D, 2A, 4B-D, 5A, 8). Micro scissors were used to remove soft tissue down to the bone, before the crania were left in 4% PFA overnight for fixation. After fixation, the cranium was rinsed in running water for 1 hour and placed in 8% hydrochloric acid (HCl) or 0.12 M EDTA in PBS for 24-48 hours for decalcification. After decalcification, the cranium was rinsed in running water for 2 hours. Rodents that did not undergo decalcification of the crania (Figure 2B-F, 3, 5B-I, 6, 7) had their brains harvested immediately after perfusion and were placed in 4% PFA overnight for fixation.

After decalcification/fixation, crania and/or brains were washed 2x in PBS for 1 hour each, then immersed in 30% and 50% ethanol for 30 minutes each, before being transferred to 70% ethanol for 24 to 72 hours at 4 °C prior to xylene and paraffin processing. The brains were embedded in paraffin, and 8-12 μm thick slices were sectioned in the coronal planes using a microtome. Slides were incubated overnight at 60 °C before sections were soaked in xylene for 20 minutes and mounted with Permount mounting medium (#SP15-100, Thermo Fisher Scientific, Waltham, MA) for light microscopy localization of AuNPs, or proceeding with immunohistochemistry for immunostaining and fluorescent microscopy co-localization of RD/TMR with meningeal fibroblast markers.

### Immunohistochemistry of RD/TMR crania

After xylene, sections were rehydrated with immersion in 100%, 95%, 70%, 50%, and 30% ethanol for 5 minutes each. After 5 minutes in deionized H2O, heat-mediated antigen retrieval was performed with 10x Diva DeCloaker (DV2004MX, BioCare Medical, Pacheco CA). Slides were cooled to room temperature before being rinsed with 1x PBS 6 times for 5 minutes each and blocked in goat serum (5% normal goat serum, 2.5% BSA, 0.5% TX-100) for 1 hour. Sections were incubated overnight at 4 °C in the following primary antibodies diluted in TBST: 1:200 dilution of anti-CRABP2 (MAB5488, EMD Millipore/Sigma Aldrich, St. Louis MO), 1:200 dilution of anti-RALDH2 (#13951-1-AP, Proteintech/Sigma Aldrich, St. Louis MO), 1:100 dilution of anti-ZO-1 (ab96587, Abcam, Cambridge MA). After incubation, sections were rinsed in PBS 6 times for 5 minutes each followed by incubation for 60 minutes at room temperature with the appropriate secondary antibodies diluted in TBST: 1:2000 dilution of goat anti-mouse IgG Alexa Fluor 488 (#A-32723, Thermo Fisher Scientific, Waltham MA), 1:2000 dilution of goat anti-rabbit IgG Alexa Fluor 647 (#A-21244, Thermo Fisher Scientific, Waltham MA). Secondary antibodies were washed off with PBS 5-6 times for 5 minutes each before incubation with a 1:500 dilution (in PBS) of 5 mg/mL DAPI for 5 minutes in the dark at room temperature. Sections were washed with PBS 3-5 times for 5 minutes each and mounted with ProLong Gold Antifade Mountant (#P36930, Thermo Fisher Scientific, Waltham MA). Confocal images were taken with the 20x and 40x (oil) lenses on a Zeiss LSM 880 Airyscan inverted two-photon microscope (Carl Zeiss Imaging, White Plains, NY).

### AuNP quantifications from light microscope images

Quantifications in Figures 2C, 3D, 5C, and 7 were obtained by first opening photomicrographs in FIJI (version 2.3.0/1.53q). Images were converted to an 8-bit image type before inverting the color. Five random areas of constant area per image were analyzed for mean density and averaged. Averaged values were compared with statistical analyses. Quantifications of dural thickness in Figure 1E were performed by measuring the straight-line distance from the outer surface of the dura to the inner surface of the dura on XRM images in FIJI.

### RD/TMR quantifications

RD/TMR tracer distribution in Figure 8 was quantified in FIJI by selecting 3 random regions from each layer of tissue that was identified in Figure 8E (CRABP2+/RALDH2+, CRABP2-/RALDH2-, CRABP2±/RALDH2-, cortex parenchyma) and measuring the mean intensity of fluorescence on the magenta/red channel in FIJI. The three regions were averaged to obtain one measurement per layer of tissue per animal. This was repeated across 4 separate animals. Averaged values were compared with statistical analyses.

#### Sample preparation for meninges analysis using gold nanoparticle-enhanced x-ray microtomography

After intracardial perfusion sacrifice 10 minutes post-1.9-nm AuNP injection and 4 hours post-15-nm AuNP injection, the entire rodent body was placed in 4% paraformaldehyde at 4° C overnight, embedded in 2% agarose, and imaged within 72 hours of sacrifice with a Zeiss Versa 520 X-ray microscope (Carl Zeiss Imaging, White Plains, NY) using a 0.4x flat panel detector. The X-ray source was tuned to 50 KV at 4W to optimally excite gold particles. Approximately 1600 projections were acquired and reconstructed, and tomograms were visualized in Zeiss XM3DViewer 1.2.8 (Carl Zeiss Imaging, White Plains, NY). Processed images were systematically reviewed in FIJI (Version 2.3.0).

#### AuNP quantifications from XRM images

For standardization, processed images from 1.9-nm and 15-nm AuNP-XRM scans were numbered by their location relative to bregma so the location of the selected images was consistent across rodents. One image per rodent (3 rodents total for 1.9-nm AuNP-XRM and 3 rodents total for 15-nm AuNP-XRM) at the level of the inferior colliculus was selected for quantification in Figures 1G-H and 4F-G and the corresponding XRM raw data file was opened in FIJI (version 2.3.0/1.53q). The mean gray intensity of three random regions within each anatomic ROI (dura, arachnoid, and pia) were calculated using FIJI/ImageJ and averaged to obtain one mean gray intensity measurement per anatomic ROI per rodent. The mean gray intensity of the agarose background was also obtained for each image and the percent change between the mean intensity of the anatomic ROIs and the mean intensity of the agarose background was calculated for standardization. A similar protocol was followed for the quantifications shown in Supplementary Figure 2, however the XRM raw data files were manually selected for each region.

### Meningeal wholemounts

Following AuNP injection, rodents were perfused with 10 mL of ice-cold PBS followed by 10 mL of 4% paraformaldehyde at 4° C. The cranium and spinal column were removed from the animal and severed at the brainstem. The cranial dura was removed by adapting a previously reported protocol [7,17]. Curved micro scissors were inserted into the foramen magnum and used to remove the top of the skull without damaging the underlying pia and brain tissue. The skullcap and brain were separately incubated in 4% PFA overnight. The dura was dissected from the bone while the leptomeninges (pia and arachnoid maters) were carefully peeled away from the brain surface.

The spinal cord was isolated from the vertebrae by using curved micro scissors to sever the pedicles bilaterally before removing the spinal cord with forceps. The dorsal half of the spinal cord was secured with forceps and its adherent dura, which appeared as a loose-hanging translucent layer, was gently peeled away and mounted on a glass slide. The dorsal leptomeninges were gently peeled away from the underlying parenchyma. This was repeated on the ventral half of the spinal cord. Wholemounts were transferred onto a glass slide, air dried, and mounted with Permount mounting medium (#SP15-100, Thermo Fisher Scientific, Waltham, MA).

### Statistical analysis

Statistical methods were not used to recalculate or predetermine sample sizes. Associations between two continuous variables were assessed using an unpaired T-test, and associations between more than two continuous variables were assessed using a one-way ANOVA with post-hoc Tukey. All tests were 2-tailed, and p-values of less than 0.05 were considered statistically significant with all p values displayed on the graphs or reported in the corresponding figure legend. All analyses were performed using Microsoft Office Excel (Version 16.36) or GraphPad Prism (Version 9.0.0)

## Results

### Transport of large CSF solutes: brain surface

We examined differential CSF handling of large CSF tracers by the cranial leptomeninges (pia and arachnoid maters) and pachymeninges (dura mater) 4 hours after intraventricular injection of 15-nm gold nanoparticles (AuNPs). Similar to the findings in previously published analyses of tracer distribution in the dura [7], 15-nm AuNP circulation within the dura, which was morphologically identified as the layer(s) immediately inferior to the skull, was restricted to the parasagittal and transverse sinus regions with only small amounts of 15-nm AuNP distribution in non-sinus regions (Figure 1B, 1C, 1D, Supplementary Figure 1A). At midline, the dura mater over the inferior colliculus was significantly thicker than the dura mater over the cortex more anteriorly and the cerebellum posteriorly (Figure 1E).

In contrast to the dura, 15-nm AuNPs distributed uniformly throughout the leptomeninges (Figure 1B, 1C, 1D). When the leptomeninges were removed, the underlying brain surface had minimal AuNP distribution, with only punctate evidence of AuNPs around the penetrating blood vessels (Supplementary Figure 1B). Leptomeningeal wholemounts of the area immediately adjacent to the middle cerebral artery (MCA) supported our histologic observations of 15-nm AuNP distribution within the leptomeningeal tissue (Supplementary Figure 1B).

We also obtained high-resolution whole-brain images using x-ray microtomography (XRM), which allowed for ex-vivo visualization of the meninges without dissection (Figure 1F). Similar to what we qualitatively observed with wholemounts and histology, XRM revealed there was significantly more 15-nm AuNP distribution within the pia mater compared to the dura mater and a trend towards increased 15-nm AuNP distribution in the arachnoid mater compared to the dura mater (Figure 1G). 15-nm AuNPs in the pia did not appear to travel inferiorly into the adjacent parenchyma (Figure 1H), as there was significantly higher 15-nm AuNP intensity in the pia compared to the parenchyma.

### Meningeal handling of large CSF solutes within the brain

On cross section, we did not observe significant differences in 15-nm AuNP distribution between non-sinus-associated dura over the dorsal surface of the brain and the falx cerebri (Figure 2A-C) [18]. We identified widespread 15-nm AuNP circulation through the leptomeninges of the perimesencephalic cisterns (Figure 2D), including the quadrigeminal, subarachnoid, ambient, and interpeduncular cisterns, and the rhinal fissure (Figure 2E). In contrast to the meninges at the surface of the brain (Figure 1H), there appeared to be CSF trafficking from the intracranial meninges into the adjacent parenchyma, as brain tissue adjacent to the perimesencephalic cisterns had a gradient of diffuse 15-nm AuNP distribution extending from the cisterns into the parenchyma (Figure 2D). 15-nm AuNPs also circulated throughout the pia mater traversing the CSF spaces intracranially, including the tela choroidea in the roof of the 3V and velum interpositum (Figure 2F). These patterns of 15-nm AuNP distribution through the tela choroidea were similar to a previously described pattern of superficial siderosis and has implications for direct communication between the ventricles outside the ependymal-lined CSF cavities [19,20]. There were no differences in the intensity of CSF solute distribution within the intracranial meninges after IVH-PHH (Supplementary Figure 2).

### Meningeal handling of large CSF solutes within the spine

We also evaluated large CSF tracer distribution within the spine meninges. Unlike the cranial dura that adheres to the cranium during dissection, the spinal dura did not adhere to the vertebrae and was dissected away from the leptomeninges (Figure 3A). The pia and arachnoid were peeled back from the underlying parenchyma in one piece (Figures 3B, 3C).

Similar to the cranial meninges, we did not observe widespread large CSF solute distribution within the spinal dura, except for diffuse AuNPs along the nerve rootlets and roots exiting the spinal cord in the lower thoracic and lumbar regions (Figure 3A). In contrast to the minimal large CSF solute handling by the dura, we observed widespread large CSF solutes circulation through the dorsal and ventral leptomeninges that outlined the structure of the median fissures and nerve roots (Figures 3B, 3C). Leptomeningeal 15-nm AuNP distribution was present in higher concentrations around the nerve roots, with additional diffuse circulation *within* the nerve roots (Figure 3D-G). On cross section of the spine, 15-nm AuNPs were seen in the arachnoid mater, pia mater, anterior median fissure, central canal, dorsal root, and root attachment zone (Figure 3H-K).

### Meningeal handling of small CSF solutes on the surface of the brain and within the brain

1.9-nm AuNP-enhanced XRM in conjunction with histology was used to evaluate small CSF tracer distribution within the dura, arachnoid, and pia. Similar to our findings with 15-nm AuNPs, non-sinus associated dura mater had significantly less 1.9-nm AuNP distribution compared to the arachnoid and pia maters 10 minutes post-intraventricular 1.9-nm AuNP injection (Figures 4A-G). While there was no overall significant difference in small CSF tracer handling between the dura over the dorsal surface of the brain and the falx cerebri, the mean intensity of 1.9-nm AuNP distribution in the falx cerebri in two out of three rodents was greater in magnitude than the mean intensity of 1.9-nm AuNP distribution in the dura over the dorsal surface of the brain in all rodents quantified (Figure 5A-C).

We also evaluated small CSF tracer handling in the intracranial leptomeninges using histology. 10 minutes after intraventricular injection of 1.9-nm AuNPs, there was notable 1.9-nm AuNP circulation in the leptomeninges of the perimesencephalic cisterns, including the ambient and interpeduncular cisterns, as well as the choroidal fissure (Figure 5D-H). This was similar to the large CSF tracer distribution patterns observed 4 hours post-15-nm AuNP injection. 1.9-nm AuNPs were also observed in the pia mater of the tela choroidea in the choroid fissure, and at the entry of the tela choroidea pia mater into the right lateral ventricle ChP (Figure 5H). This was akin to the 15-nm AuNP distribution seen in the dorsal 3V and velum interpositum. There was also 1.9-nm AuNP circulation in the pia mater in the folia of the cerebellum (Figure 5I). No significant differences in small CSF tracer distribution within the intracranial meninges were observed after IVH-PHH (Supplementary Figure 2).

### Meningeal handling of small CSF solutes within the spine

Similar to what we observed with large CSF solutes, there was broad small CSF solute distribution through the spinal leptomeninges 10 minutes after intraventricular injection, but not within the dura mater (Figure 6A). On cross section, 1.9-nm AuNPs were concentrated in the anterior median fissure, ventral horn, dorsal root, and root attachment zone (Figure 6A, 6B). 1.9-nm AuNPs were also seen within the invaginations around penetrating blood vessels (Figure 6A). Notably, 1.9-nm AuNPs were present in the cell bodies of the dorsal root ganglia (Figure 6A, 6C); the distribution among the cell bodies was not homogenous, with some cell bodies taking up more AuNPs than others (Figure 6C).

### CSF solute handling by the choroid plexus

Given the close developmental relationship between the tela choroidea and ChP, we evaluated ChP handling of large and small CSF solutes. Large solutes were present in the lateral ventricle (LV), anterior third ventricle (3V), and posterior 3V ChPs (Figure 7A) 4 hours after intraventricular injection. There was significantly more apical than luminal 15-nm AuNP distribution in the anterior and posterior 3V ChPs, but no difference in the LV ChP (Figure 7B-D). When comparing 15-nm AuNP distribution between the LV, anterior 3V, and posterior 3V ChP, there was significantly more 15-nm AuNP distribution in the posterior 3V ChP compared to the anterior 3V ChP 4 hours post-injection (Figure 7E, 7F). The ChP epithelial cells did not take up 15-nm AuNPs into their cell bodies.

In contrast to large CSF solutes, there was significantly decreased small CSF solute (1.9-nm) distribution on the apical side in the 3V and 4V ChPs compared to the luminal side 10 minutes post-injection, but not in the lateral ventricle ChP (Figure 7G-J). There was also increased 1.9- nm AuNP distribution on the luminal side of the 3V and 4V ChP compared to the LV ChP (Figure 7K), however no difference was observed on the apical side between the three locations (Figure 7L).

### Fluorescent CSF tracer colocalization with meningeal fibroblast markers

To verify the identity of the meningeal layers interacting with CSF tracer, we injected 3kDa fluorescent CSF tracer RD/TMR into the right lateral ventricle to allow for tracer co-localization with meningeal fibroblast markers including the arachnoid fibroblast markers retinaldehyde dehydrogenase 2 (RALDH2) [16], the arachnoid and dural fibroblast marker cellular retinoic acid binding protein 2 (CRABP2) [16], and the tight junction protein zonula occludens 1 (ZO-1). RD/TMR co-localized with layers of cells that were strongly positive for CRABP2 and RALDH2, and another layer that was CRABP2-/RALDH2- located superior to the brain parenchyma but inferior to the CRABP2+/RALDH2+ layer (Figure 8A-C). Both of these layers were inferior to a CRABP2+/ZO- 1 layer, suggesting they represent the arachnoid and pia maters. Within cross sections of the falx cerebri, there were additional layers of tissue medial to the strongly CRABP2+ layer that did not have RD/TMR distribution that likely represented the dura mater (Figures 8A, 8B), which is anatomically medial to the arachnoid, pia, and parenchyma at midline. The CRABP2+/RALDH2+ (arachnoid) and CRABP2-/RALDH2- (pia) layers had significantly more RD/TMR CSF tracer than the more superior CRABP2±/RALDH2- layer (dura) (Figure 8E), which is consistent with our observations with AuNPs that relied on morphologic identification of the meningeal layers.

## Discussion

Recent studies have brought new perspective to longstanding theories of CSF circulation that predominantly implicate CSF absorption at arachnoid granulations [1,6,7]. Despite the reinvigorated focus on the meninges, the CSF distribution through the entire meninges, particularly the developing intracranial leptomeninges (pia and arachnoid maters) and spine meninges is not well known [21]. In this study, we report the distribution of large and small CSF tracers within the meningeal and ventricular networks. We show that after injection into the lateral ventricle, small and large CSF tracers distribute through the cranial and spinal leptomeninges, notably in the perimesencephalic cisterns and tela choroidea. In the choroid plexus, small tracers were localized to the lumen, with differential distribution between the ventricles; the 4V and 3V choroid plexus had significantly more tracer than the lateral ventricle. Large tracers were localized to the apical choroid plexus with significantly more tracer in the posterior 3V compared to the anterior 3V.

By closely analyzing the macroscopic distribution of CSF tracers through the meninges and choroid plexus, we hope to call attention to the role of structures outside the arachnoid granulations and dura in CSF solute distribution, especially during the neonatal time period. The differential handling of CSF tracers by the dura and leptomeninges may suggest that the different layers of the meninges have distinct roles in CSF handling. The regional differences in small CSF tracer handing between the dura over the surface of the brain and the dura within the interhemispheric invagination (ie. the falx cerebri) in two out of three rodents suggest that the slightly higher amounts of small CSF tracer within the falx cerebri may be secondary to the falx cerebri playing an role in mediating CSF-dural sinus interactions. Specifically, the falx cerebri is in unique proximity to the superior sagittal sinus. The inferior sagittal sinus courses through the lower boundary of the falx cerebri, suggesting the falx cerebri may provide a local environment extending down the depth of the cortex in which CSF, proteins, and molecules from the ISF and brain parenchyma are able to be trafficked into the parasagittal meningeal lymphatics to allow for ongoing drainage and immune monitoring, particularly in older animals with functional meningeal lymphatics [7,22–27]. Further investigations should be done into the nature and rate of transport of CSF, CSF tracer, protein, and other molecules into the falx compared to other dural regions.

The perimesencephalic cisterns surround the midbrain and include the ambient, crural, interpeduncular, and quadrigeminal cisterns [28]. The location of these cisterns is remarkable in that they separate two functionally-distinct regions – the hippocampus and the midbrain. This, in conjunction with the results of our present study showing diffuse 15-nm AuNP movement into only the hippocampal side of the quadrigeminal, subarachnoid, and ambient cisterns, but not the midbrain, suggest that these cisterns may serve to facilitate a regionalized CSF-brain parenchyma cross talk [29–32]. Alternatively, their location in between the lateral and third ventricles may serve as a potential route for inter-ventricular CSF transport and communication [20]. While we did not see any significant differences in tracer distribution within the meninges of these cisterns after IVH-PHH, it is possible there are functional differences allowing increased/decreased CSF solute transport from the cisterns into the adjacent parenchyma.

In addition to the perimesencephalic cisterns, one particularly intriguing yet understudied intracranial meningeal structure is the tela choroidea, a thin region of pia mater adherent to the underlying fourth ventricle ependyma [33–38]. In the third ventricle, the tela choroidea forms the roof of the ventricle and superimposes upon itself to form the velum interpositum, which lies between the internal cerebral veins and contains cerebrospinal fluid [33,35,37,39,40]. The underlying ependyma of the tela choroidea also gives rise to the ChP, forming a continuous pial structure from its initiation near the cerebellum to the choroidal fissure in the medial wall of the lateral ventricles. The tela choroidea has been purported to have a role in allowing CSF to be recirculated into the ventricular system but has otherwise been sparsely studied in the context of CSF circulation [19,41,42]. In this study, we report that 1.9-nm AuNPs were found predominantly in the luminal side of the 3V and 4V choroid plexus as soon as 10 minutes post-injection. There was significantly less luminal 1.9-nm AuNP in the LV choroid plexus compared to the 3V and 4V. In addition, there was significantly less apical, compared to luminal 1.9-nm AuNP distribution in all of the ventricles, suggesting that the 1.9-nm AuNPs were not being shuttled across the choroid plexus epithelial cells into the lumen, but rather through direct transportation along the luminal pia mater. It is possible that the first point of 1.9-nm AuNP entry to the choroid plexus is through the tela choroidea of the fourth ventricle, after which it traveled anteriorly along the tela choroidea to reach the roof of the third ventricle and eventually the choroidal fissure and lateral ventricle choroid plexus. A previous study on experimental hydrocephalus in primates and dogs posited that CSF traveled to the ventricles from the cisterns via the tela choroidea [42,43]. Other studies have indicated that the CSF may be transported across a normally functioning tela choroidea into the choroid plexus and ventricles as a regular aspect of brain physiology [19,41]. The results of our present study support these findings and build upon previous studies implicating the choroid plexus to be involved in direct CSF solute transport [44,45].

Our investigations were conducted ex vivo, which may impact the structure and presence of CSF spaces such as the subarachnoid space. Similarly, while decalcification with EDTA and HCl allowed for visualization of the skull, meninges, and brain in approximation with each other, it resulted in disfiguration of the skull over the dorsal convexities. Additionally, our investigation was based on rodents, animals in which the presence, morphology, and function of arachnoid villi have been debated. Finally, the animal we used for this study is lissencephalic, which may affect the leptomeningeal distribution of CSF tracers in the cerebrum because the pia mater follows the invaginations produced in animals with gyri and sulci. In the cerebellum, which retains its sulci and gyri in the rodent brain, there was an abundance of CSF tracer in the pia mater invaginations into the folia. Repeating these experiments in organisms with gyri and sulci, such as pigs or ferrets, may show increased leptomeningeal tracer in the leptomeninges over the cerebral cortex.

## Conclusion

We present a CNS-wide map of meningeal handling of large and small CSF solutes in the neonatal brain and spine. Intracranial leptomeninges, particularly the tela choroidea in the fourth ventricle and at the roof of the third ventricle, are an important area for CSF circulation and future studies should be performed to characterize mechanisms of CSF transport along these areas. Differential choroid plexus distribution of tracers by ventricle and between the luminal and apical surfaces suggests different roles of these regions in facilitating CSF flow. Finally, in contrast to prior reports of dural CSF solute transport in adult animals, we show that meningeal handling of both large and small CSF solutes is mediated by the pia and arachnoid in young animals.

## Abbreviations

AuNP: gold nanoparticle
ChP: choroid plexus
CNS: central nervous system
CSF: cerebrospinal fluid
IVH: intraventricular hemorrhage
PHH: posthemorrhagic hydrocephalus
RD/TMR: red dextran tetramethylrhodamine
SAS: subarachnoid space
3V: third ventricle
4V: fourth ventricle

## Acknowledgements

This work was funded by the National Institutes of Health (R01 NS110793 to J.M.S.), K12 Neurosurgeon Research Career Development Program (J.M.S), and the Hydrocephalus Association (J.M.S), McDonnell Center for Systems Neuroscience at Washington University in St. Louis (J.M.S.), the Children’s Discovery Institute (J.M.S.), and the Washington University in St. Louis Center for Cellular Imaging also known as WUCCI (J.M.S.). XRM imaging was performed with a Zeiss Xradia 520 X-ray Microscope at WUCCI, which is supported by Washington University in St. Louis School of Medicine and the NIH (OD021694). Figures were created with BioRender.com.

## Data availability statement

The datasets generated during and/or analysed during the current study are available from the corresponding author on reasonable request.

