## Supplementary Figure 1 for "Meningeal CSF transport is primarily mediated by the arachnoid and pia maters during development"

**Supplementary Figures**

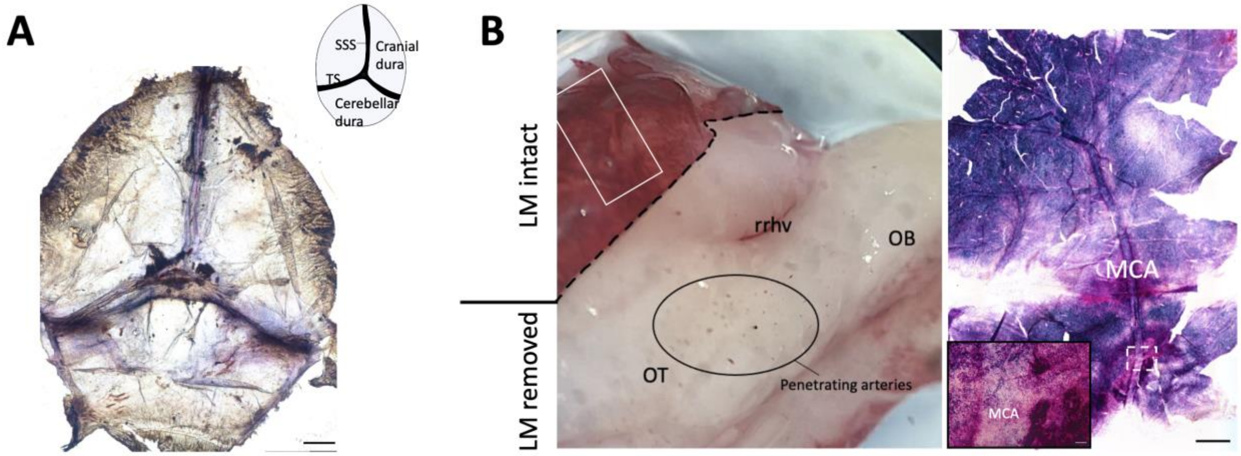


**Supplementary Figure 1. A**, Representative image of 15-nm gold nanoparticle (AuNP) distribution in the dura 4 hours after intraventricular injection in P7 rodents. Inset, schematic demonstrating anatomy of whole-mount dissection of the rat brain dura. SSS, superior sagittal sinus; TS, transverse sinus. scalebar = 1 mm. **B**, Dissection microscope image of harvested P7 rat brain 4 hours after 15-nm AuNP injection with the leptomeninges (pia-arachnoid) peeled back from the base of the brain. Solid white box indicates the location that the leptomeningeal wholemount (right) showing the middle cerebral artery (MCA) is taken from. Inset, higher magnification image of the MCA on the LM wholemount. olfactory tubercle; rrhv, rostral rhinal vein; OB, olfactory bulb. scalebar = 1 mm, inset scalebar = 50 µm
