## Supplementary Figure 2 for "Meningeal CSF transport is primarily mediated by the arachnoid and pia maters during development"

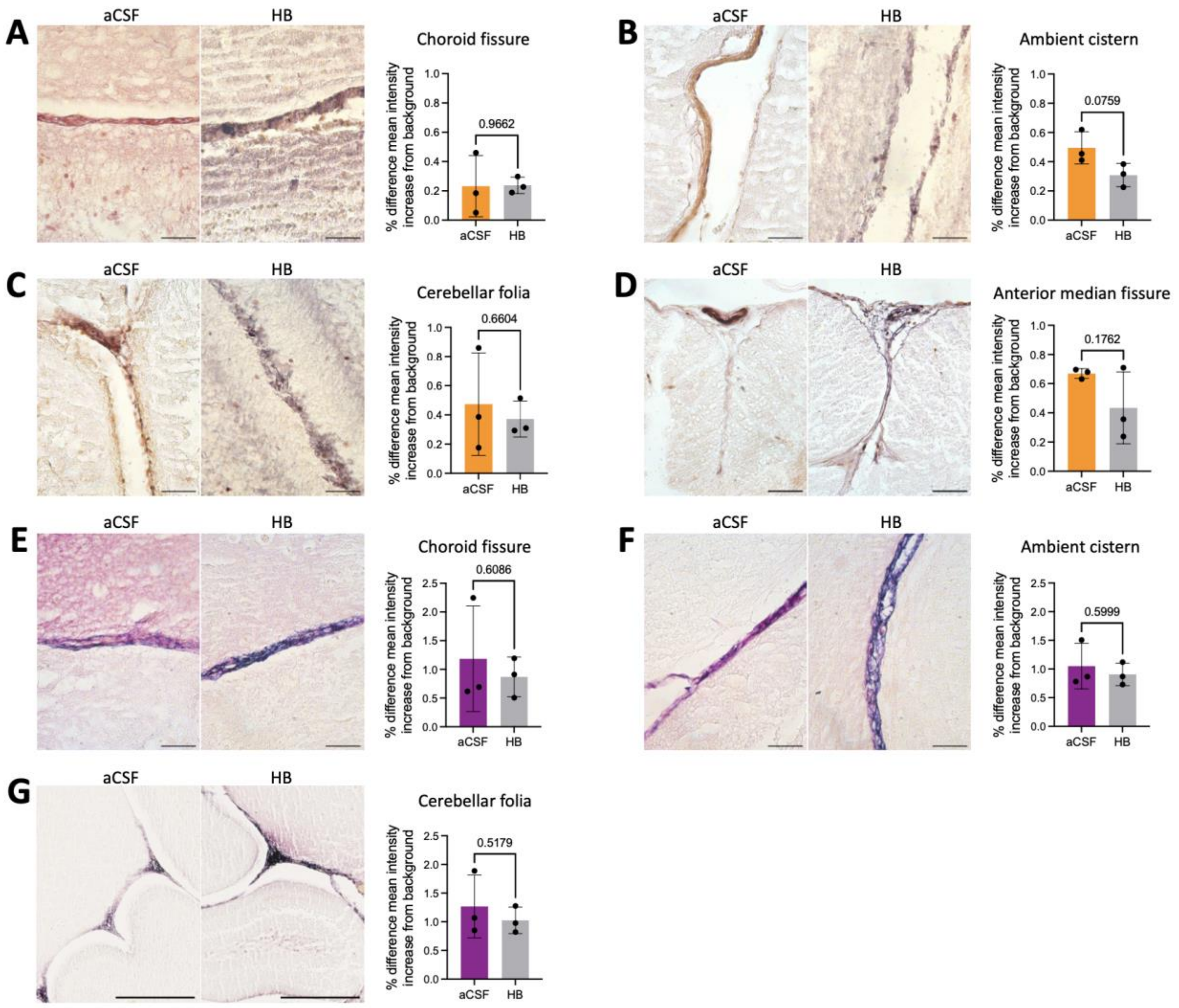


**Supplementary Figure 2.** Representative images of 1.9-nm gold nanoparticle (AuNP) (A-D) and 15-nm AuNP (E-G) distribution in neonatal rodents with intraventricular hemorrhage-posthemorrhagic hydrocephalus (IVH, induced through intraventricular hemoglobin (HB)) compared to aCSF controls 72 hours post-IVH induction. **A-D**, 1.9-nm AuNP distribution in the choroidal fissure (A), ambient cistern (B), cerebellar folia (C), and anterior median fissure of the spinal cord (D) 10 minutes post-1.9-nm AuNP injection into the right lateral ventricle of IVH and control neonatal rodents. A-D scalebars = 50 µm. **E-G**, 15-nm AuNP distribution in the choroidal fissure (E), ambient cistern (F), and cerebellar folia (G) 4 hours post-15-nm AuNP injection into the right lateral ventricle of IVH and control neonates. E-F scalebars = 50 µm, G scalebars = 250 µm. Quantifications are also shown as mean ± SD with unpaired two-tailed t tests, n = 3 animals per group.
