## Supplementary Information for "Meningeal CSF transport is primarily mediated by the arachnoid and pia maters during development"

**Supplementary methods**

*Meningeal wholemounts.* Following AuNp injection, animals were perfused with 10 mL 4% paraformaldehyde at 4º C and 10 mL ice-cold PBS and placed dorsal side up. A single midline cut was made from the nose to the tail and the skin retracted to expose the skull, muscle and underlying tissues. Spinal column excision was performed by making an incision in the region of the lower back/femurs 0.5 cm lateral to the column, and cutting the back musculature up towards the head, cutting through the hip joint, ribs, and shoulder joint to remove the arm and leg, with care taken to keep the central nervous system structures intact. This was repeated on the contralateral side. The viscera attached on the ventral side of the spinal column was cut to free the spinal column and head from the rest of the body. Using curved microsurgical scissors, with the tips pointing away from the brain, the posterior atlanto-occipital membrane and cisterna magna were cut, exposing the foramen magnum. Sliding the curved scissors into the foramen magnum, the skull was cut counterclockwise, superior to the posttympanic hook and the zygomatic process of the frontal bone and around the anterior-most aspect of the frontal bone to remove the frontal, parietal, interparietal, and occipital bones and the adherent underlying dura in one piece. The curved scissors were then held parallel to the table, with tips pointing away from the spinal cord, and used to sever the pedicles of the vertebra to remove the vertebral bodies and expose the dorsal side of spinal cord. The scissors were rotated 90 degrees and used to cut the vertebral lamina and sever the dorsal roots as close to the cord as possible. The brain and spinal cord tissue were carefully separated from the remaining bone in one piece, taking care to keep the dura and arachnoid intact, and left in 4% PFA overnight at 4 degrees.

The following day, the tissue was placed in a petri dish with ice-cold PBS under a dissecting microscope. The brain and spinal cord were severed. Using curved forceps to gently secure the spinal cord, microsurgical scissors were used to cut the dura and leptomeninges along the length of both sides of the spinal cord. After orienting the spinal cord with the dorsal side facing up, the dura, which appeared as a loose, translucent layer, was gently peeled away from the leptomeninges and parenchyma in one piece. Next, the leptomeninges, which appeared as a spongy layer adherent to the underlying parenchyma, was gently peeled away from the underlying parenchyma. This was repeated on the ventral side. The cranial leptomeninges were removed in three pieces, one from each cerebral hemisphere, and one from the cerebellum. To remove the leptomeninges from the cerebral hemispheres, a midline incision was made in the leptomeninges on the base of the brain. Using curved forceps, the leptomeninges were carefully peeled from the incision moving up towards the longitudinal fissure and removed as one large piece. The cerebellar leptomeninges were removed by using the curved forceps to detach the leptomeninges from the superior colliculi and gently peeling tissue from the folia, moving down towards the foramen magnum.

Dura and pia-arachnoid wholemounts were transferred onto a glass slide, air dried, and mounted with Permount mounting medium (#SP15-100, Thermo Fisher Scientific, Waltham, MA) for imaging with light microscopy.
